# Atheroprone shear stress stimulates noxious endothelial extracellular vesicle uptake by MCAM and PECAM-1 cell adhesion molecules

**DOI:** 10.1101/2022.12.31.522373

**Authors:** Pierre-Michaël Coly, Shruti Chatterjee, Fariza Mezine, Christelle El Jekmek, Cécile Devue, Thomas Nipoti, Maribel Lara Corona, Florent Dingli, Damarys Loew, Guillaume van Niel, Chantal M. Boulanger

## Abstract

Atherosclerotic lesions mainly form in arterial areas exposed to low shear stress (LSS), where endothelial cells express a senescent and inflammatory phenotype. Conversely, high shear stress (HSS) has atheroprotective effects on the endothelium. Endothelial cell-derived extracellular vesicles have been shown to regulate inflammation, senescence and angiogenesis and therefore play a crucial role in vascular homeostasis and disease. While previous studies have shown links between hemodynamic forces and extracellular vesicle release, the exact consequences of shear stress on the release and uptake of endothelial EVs remains elusive. Our aim is therefore to decipher the interplay between these processes in endothelial cells exposed to atheroprone or atheroprotective shear stress.

Confluent human umbilical vein endothelial cells (HUVEC) were exposed to either LSS or HSS for 24 hours. Large and small EVs were isolated from conditioned medium by sequential centrifugation and size exclusion chromatography. They were characterized by TEM, Western blot analysis of EV markers, tunable resistive pulse sensing, flow cytometry and proteomics. Uptake experiments were performed using fluorescently-labeled EVs and differences between groups were assessed by flow cytometry and confocal microscopy.

We found that levels of large and small EVs in HUVEC conditioned media were fifty and five times higher in HSS than in LSS conditions, respectively. *In vivo* and *in vitro* uptake experiments revealed greater EV incorporation by cells exposed to LSS conditions compared to HSS. Additionally, endothelial LSS-EVs appeared to have a greater affinity for HUVECs than HSS-EVs or EVs derived from platelets, red blood cells, granulocytes and peripheral blood mononuclear cells. Proteomic analysis revealed that LSS-EVs were enriched in adhesion proteins such as PECAM1, MCAM, which were involved in EV uptake by endothelial cells. LSS-EVs also carried mitochondrial material, which may be involved in elevating reactive oxygen species levels in recipient cells.

These findings suggest that endothelial shear stress has a significant impact during EV biogenesis and uptake. Given the major role of EVs and shear stress in vascular health, deciphering the relation between these processes may yield innovative strategies for the early detection and treatment of endothelial dysfunction.

## Introduction

Hemodynamic forces are a major determinant of atherosclerotic plaque formation. Shear stress, the mechanical force exerted by the blood flow on the vascular wall, has a profound impact on the phenotype of endothelial cells (Chiu & Chien, 2011; Gimbrone & García-Cardeña, 2013; Hahn & Schwartz, 2009). Plaques preferentially form in regions where the blood flow is turbulent and oscillatory, such as arterial bifurcations or curvatures. Here, endothelial cells are exposed to low shear stress (LSS), which favor apoptosis, senescence and inflammation. Conversely, regions of laminar blood flow are exposed to high shear stress (HSS) and are relatively protected from atherosclerotic lesions (Davies et al., 2013; Tricot et al., 2000; Warboys et al., 2014). Among the affected pathways, our group and others demonstrated that LSS deeply impacts the endothelial endolysosomal system, namely through reduced lysosomal acidity and perturbed autophagic flux (Chung et al., 2022; Liu et al., 2015; Vion et al., 2017).

Extracellular vesicles (EVs) are submicron membrane particles released by all cell types *via* diverse mechanisms. The two main EV subpopulations are exosomes (50-150nm) and microvesicles (100-1000nm). The former is created by inward budding of the endosomal membrane, thereby forming intraluminal vesicles, and secreted when multivesicular endosomes fuse with the plasma membrane. Microvesicles, on the other hand, are generated by outward budding of the plasma membrane (van Niel et al., 2018). Long considered as only a means of eliminating cellular waste, EVs are now believed to be key players in cellular communication, as they can carry biological information (protein, lipids, nucleic acids) between cells (van Niel et al., 2022). Being in direct contact with the blood flow means that endothelial cells can easily release EVs into the circulation, this makes them the second contributor of circulating EVs behind platelets (Mathiesen et al., 2021). The endothelium also interacts with circulating EVs, as they have been seen rolling, arresting and accumulating on the endothelial cell walls *in vivo* (Hyenne et al., 2019; Verweij et al., 2019). The composition and subsequent effect of EVs seems to be largely dependent on the state of the parental cell. In that’s sense, EVs are involved in several physiopathological processes, many of which underlie the progression of atherosclerosis. EVs derived from injured of inflamed endothelial cells have been shown to carry specific microRNAs, cytokines or adhesion proteins, and have been implicated in monocyte activation, endothelial permeability and smooth muscle cell proliferation (Coly & Boulanger, 2022). While it is known that shear stress is a major determinant of EV production (Vion et al., 2013), we have relatively little data on the impact of these forces of the full composition of endothelial EVs. Similarly, we know little on how the level of shear stress affects EV endocytosis by endothelial cells, and the consequences on the endothelial phenotype. Interestingly, recent data have shown that the process of EV release can intersect with autophagy, be it by the formation of secretory amphisomes upon fusion of autophagosomes with multivesicular endosomes, or the fusion of mature autophagosomes with the plasma membrane (van Niel et al., 2022). These mechanisms expand the variety of EV subpopulations and result in the release of heterogenous content into the extracellular space, including digestion material.

By impacting the endolysosomal network we hypothesize that shear stress can alter the endocytosis of EVs by endothelial cells, as well as the composition of secreted EVs. In this study, we found that endothelial cells exposed to atheroprone LSS incorporated more EVs than cells exposed to HSS *via* macropinocytosis and a clathrin-dependent pathway. Endothelial cells exposed to LSS released EVs that were enriched in adhesion and mitochondrial proteins. Functionally, this resulted in LSS-EVs being more readily endocytosed than HSS-EVs, as well as having a pro-oxidative effect on recipient cells.

## Results

### Shear stress affects the concentration of EVs found in endothelial cell conditioned media

HUVECs were exposed to HSS or LSS conditions for 24 h. Large (lEVs) and small (sEVs) EVs were then isolated from the conditioned media by differential centrifugation followed by size exclusion chromatography, and their morphology was observed by electron microscopy (Fig. 1A). Quantification by tunable resistive pulse sensing revealed that sEVs and lEVs had a mode diameter of approximately 100 nm and 230 nm respectively, with no significant difference between HSS and LSS EVs. In terms of concentration, we found substantially more EVs in the conditioned medium of cells exposed to HSS, compared to LSS (Fig. 1B, 1C and 1D). Finally, Western blot analysis revealed that EVs harbored the transmembrane proteins CD63, CD9 and the cytosolic protein HSC70 (Fig. 1E).

**Figure 1:**
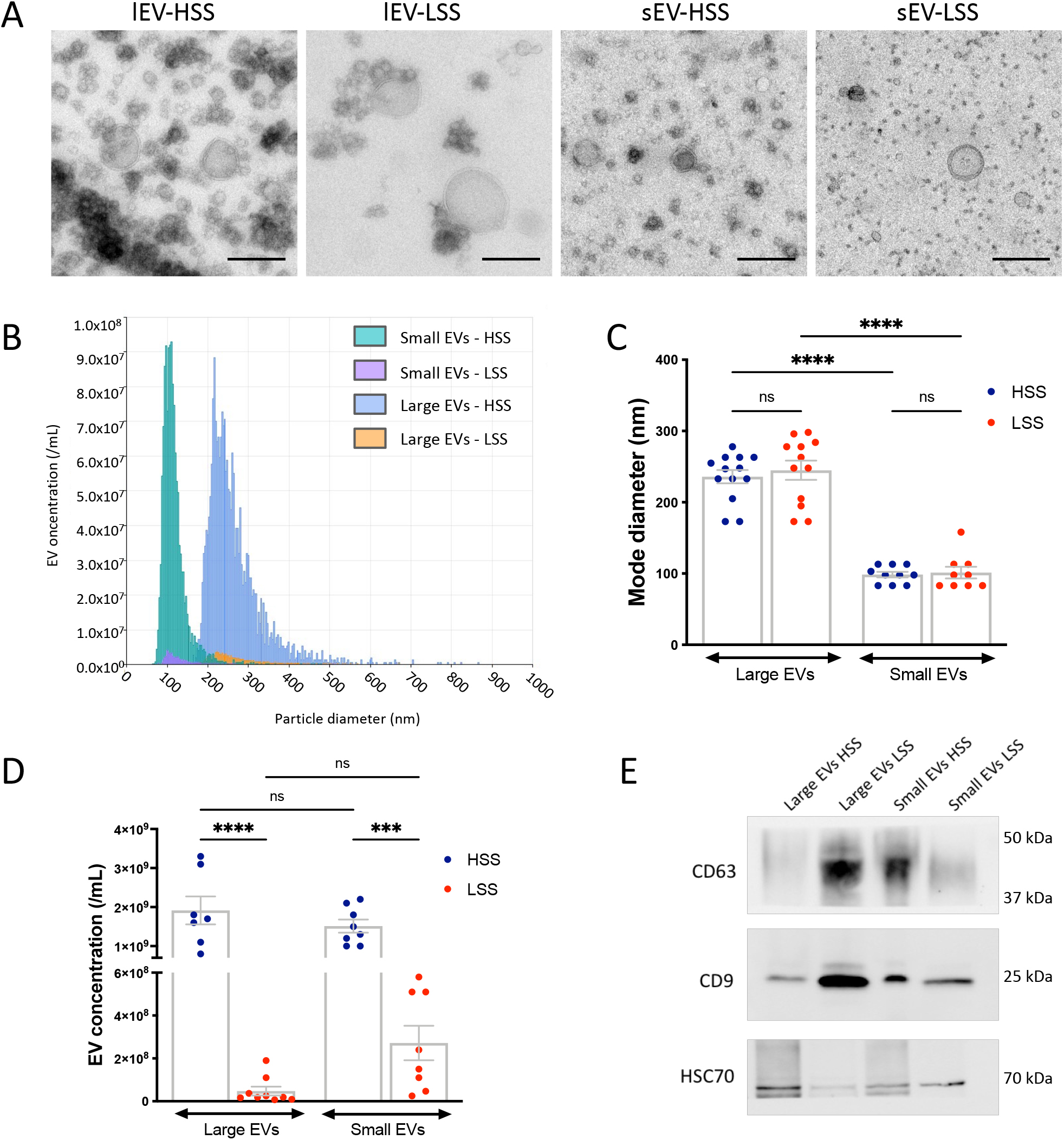
Characterization of endothelial EVs produced under HSS or LSS conditions. **(A)** HUVECs were exposed to HSS or LSS for 24h. Large (lEVs) and small (sEVs) EVs were isolated by differential centrifugation and size exclusion chromatography and observed by transmission electron microscopy. Scale bar = 200 nm. **(B)** Representative size distribution analysis by tunable resistive pulse sensing (TRPS). **(C)** EV mode diameter determined by TRPS. **(D)** EV concentration determined by TRPS. **(E)** Representative Western blot for CD63, CD9 and HSC70 in lEVs and sEVs produced under HSS or LSS conditions. Data represent means ± SEM of 7-13 independent experiments. ***P<0.001, ****P<0.0001; One-way Anova.

### EVs produced under LSS conditions are more readily taken up than EVs produced under HSS

To assess if endothelial EVs could be taken up by endothelial cells *in vivo*, we labeled murine endothelial cell-derived EVs (SVEV4-10) with the fluorescent lipophilic dye Membright as described in Hyenne et al., 2019. Mice were injected with labeled-EVs and sacrificed after 30 minutes to assess EV incorporation into vascular endothelial cells. Confocal imaging of mouse aortas revealed that HSS-lEVs had little retention in endothelial cells, as endothelial fluorescence was bellow detection levels. Conversely LSS-lEVs were visible in the mouse aorta following injection and seemed to preferentially accumulate in the inner aortic arch (region of LSS) when compared to the thoracic aorta (region of HSS; Fig. 2A). We then performed *in vitro* EV-uptake assays using fluorescently-labeled endothelial EVs produced under HSS or LSS conditions. We deposited these EVs on HUVECs that had been exposed to HSS or LSS for 24 h, and assessed the rate of EV accumulation by confocal microscopy and flow cytometry. Confocal imaging revealed that HUVECs exposed to LSS took up more fluorescently-labeled EVs than HUVECs exposed to HSS (Fig. 2B and 2C), thus corroborating our *in vivo* observations (Fig. 2A). Here also, we found that LSS-lEVs were more readily detectable in endothelial cells, suggesting greater uptake of these vesicles compared to HSS-lEVs (Fig. 2D). sEV uptake was bellow detection levels by this method. Similar data were obtained when analyzing cells by flow cytometry. LSS-EVs were detectable in 10-20 % of cells, while HSS-EVs were detectable in less than 5% of cells (Fig. 2E). Further examination by imaging flow cytometry showed that the fluorescent EV pattern often appeared to be located within the cytosol, rather than only at the cell surface, which suggests that these EVs are being endocytosed by HUVECs (Fig. 2F). To ensure that this was the case, we targeted several endocytic pathways previously described for EV uptake, i.e., macropinocytosis, clathrin-mediated endocytosis and caveolin-mediated endocytosis using pharmacological inhibitors: amiloride, chlorpromazine and nystatin respectively. We found that perturbing macropinocytosis and clathrin-mediated endocytosis reduced the EV signal detected in cells, which indicates that these pathways are necessary for endothelial EV uptake by HUVECs (Fig. 2G). We next sought to compare the uptake levels of endothelial EVs to plasma-derived EVs, as well as EVs isolated from circulating cells: peripheral blood mononuclear cells, neutrophils, platelets, and red blood cells. While all EV types seemed to be taken up by HUVECs, endothelial EVs produced under LSS conditions displayed the highest rate of uptake, suggesting a higher affinity of endothelial cells for these vesicles (Fig. 2H).

**Figure 2:**
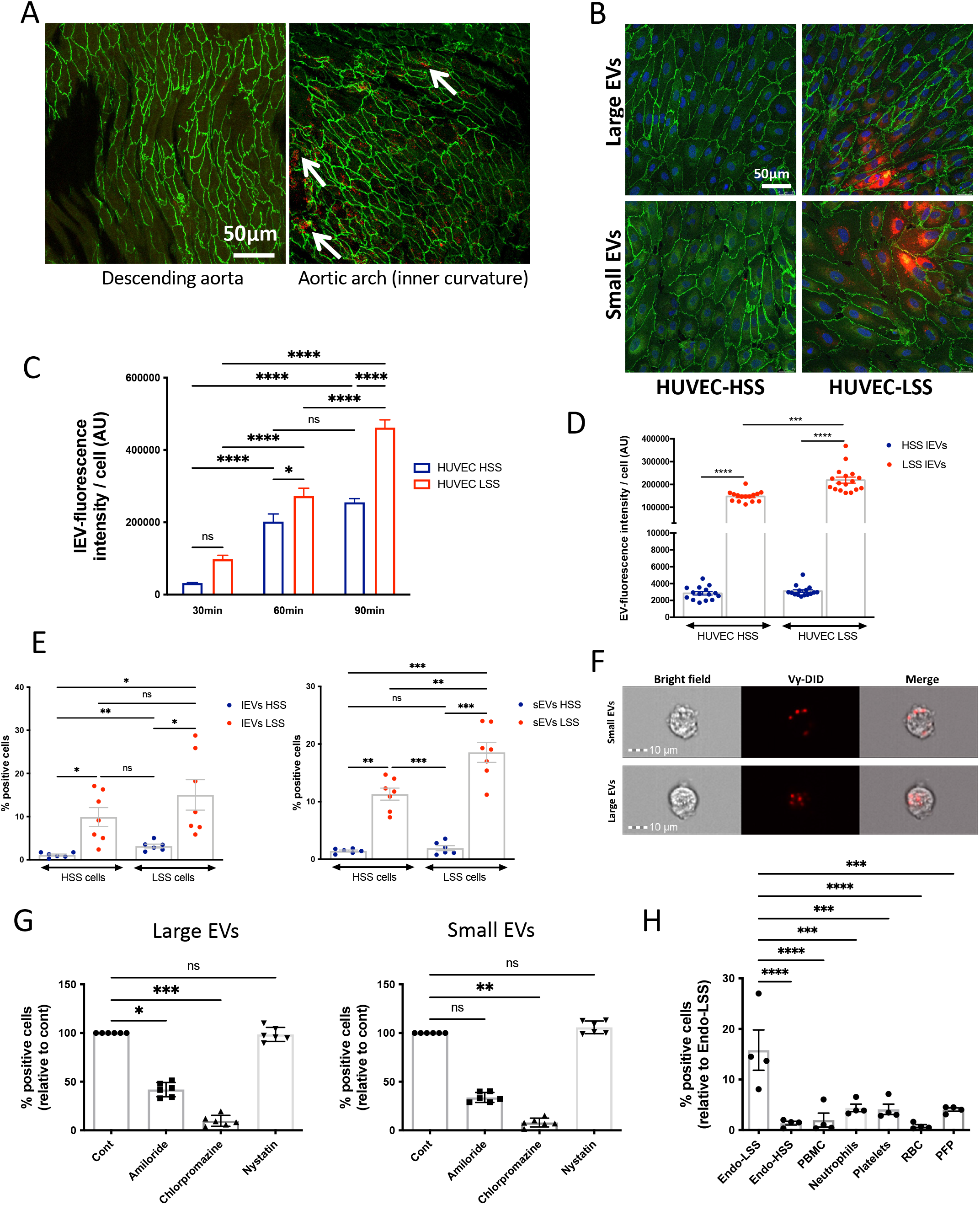
LSS-EVs are more readily taken up than HSS-EVs. **(A)** Fluorescently-labeled SVEC4-10 LSS-lEVs (red) were injected into the mouse bloodstream. Animals were sacrificed after 30 min and aortas were harvested. Endothelial cells (green; Cadherin-5 staining) were imaged by confocal microscopy. **(B)** Fluorescently-labeled HUVEC LSS-lEVs (red) were incubated for 90 min with HUVECs exposed to HSS or LSS conditions for 24 h. Cells were fixed, stained for Cadherin-5 (green) and imaged by confocal microscopy. **(C)** Fluorescently-labeled HUVEC-derived LSS-lEVs were incubated for varying periods min with HUVECs exposed to HSS (blue) or LSS (red) conditions for 24 h. Representative quantifications of EV signal per cell from confocal images captured at different time points (15 fields per condition, *P<0.05, ****P<0.0001, One-way Anova). **(D)** Fluorescently-labeled HUVEC-derived HSS-lEVs (blue) or LSS-lEVs (red) were incubated for 90 min with HUVECs exposed to HSS or LSS conditions for 24 h. Representative quantifications of EV signal per cell from confocal images (15-17 fields per condition, ***P<0.001, ****P<0.0001, One-way Anova). **(E)** Fluorescently-labeled HUVEC-derived HSS-(blue) or LSS-(red) lEVs (left) or sEVs (right) were incubated for 90 min with HUVECs exposed to HSS or LSS conditions for 24 h. % of cells positive for EV signal was analyzed by flow cytometry. Data represent means ± SEM of 7 independent experiments, *P<0.05, **P<0.01, ***P<0.001, One-way Anova. **(F)** Fluorescently-labeled HUVEC-derived LSS lEVs or sEVs were incubated with HUVECs exposed to LSS conditions for 24 h. Cells were and visualized by image flow cytometry. **(G)** Fluorescently-labeled HUVEC-derived LSS lEVs (left) or sEVs (right) were incubated for 90 min, in the presence of endocytosis inhibitors, with HUVECs exposed to LSS conditions for 24 h. % of cells positive for EV signal, relative to Cont, was analyzed by flow cytometry. Data represent means ± SEM, relative to Cont, of 6 independent experiments, *P<0.05, **P<0.01, *** P<0.001, Friedman test. **(H)** Fluorescently-labeled lEVs derived from HUVECs exposed to LSS (Endo-LSS) or HSS (Endo-LSS), peripheral blood mononuclear cells (PMBC), neutrophils, platelets, red blood cells (RBC) or platelet free plasma (PFP) were incubated for 90 min with HUVECs exposed to LSS conditions for 24 h. % of cells positive for EV signal was analyzed by flow cytometry. Data represent means ± SEM of 4 independent experiments, ***P<0.001, ****P<0.0001, One-way Anova.

### Quantitative proteomic analysis of large and small EVs produced under HSS and LSS conditions

To shed light on the mechanisms underlying the higher uptake of LSS-EVs, we performed a proteomic analysis of large and small EVs produced under HSS and LSS conditions, from five biological replicates. We identified a variety of proteins traditionally found in mammalian EVs, including tetraspanins and heat shock proteins (Supp. Table 1). Furthermore, Gene Ontology (GO) term enrichment analysis of these proteins showed terms relating to extracellular vesicles (Supp. Fig. 1). Qualitative analysis, evaluating the presence of proteins identified by at least three distinct peptides, indicated that the majority of proteins were common to all four groups (Fig. 3A). Each group also contained a unique set of proteins, with LSS EV sets being more substantial. GO term enrichment analysis of unique proteins revealed accumulation of cell adhesion proteins in LSS-lEVs and ribosome proteins in LSS-sEVs.

**Figure 3:**
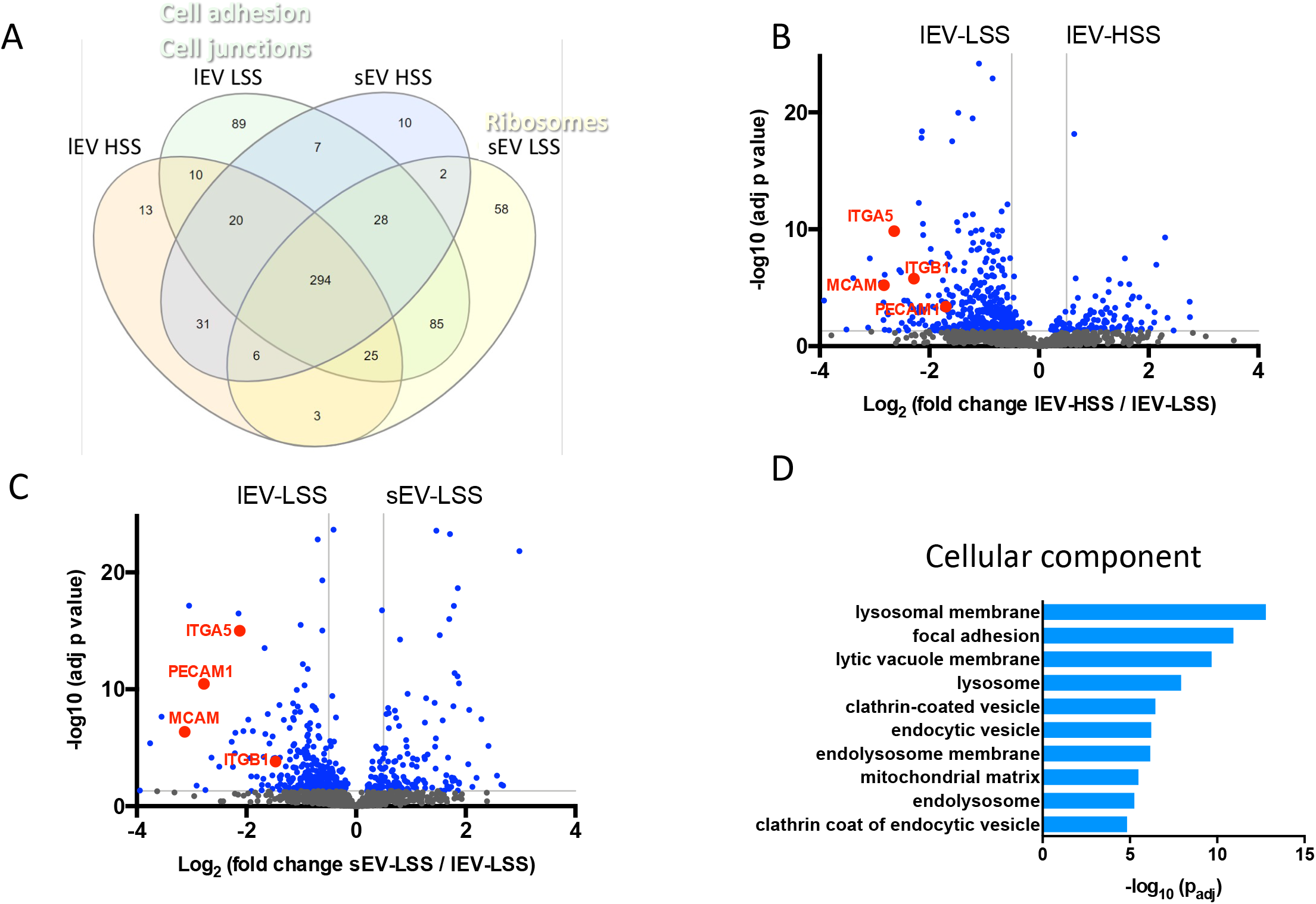
Qualitative and quantitative proteomic analysis of endothelial EVs produced under HSS and LSS. **(A)** Venn diagram showing the distribution of proteins qualitatively identified in each group by at least three peptides in five biological replicates. Quantitative analysis of proteins present in HSS-lEVs **(B)** or LSS-sEVs **(C)** compared LSS-lEVs is shown as Volcano plots. X axis = log_2_ (fold change), Y axis = −log10 (adj P value). The horizontal grey line indicates P value = 0.05, vertical grey lines indicate absolute fold-change = 2. **(D)** Gene Ontology, cellular component terms associated with proteins enriched in LSS-lEVs.

As most proteins were common between all groups, we then performed a quantitative comparison of their amounts in LSS-lEVs vs HSS-lEVs (Fig. 3B) or LSS-lEVs vs LSS-sEVs (Fig. 3C). GO term classification of proteins significantly (P < 0,05) more abundant (≥ 2) in LSS-lEVs compared to HSS-lEVs or LSS-sEVs, highlighted “extracellular space”, “cell junction” “focal adhesion” which suggests that LSS-lEVs are equipped with a higher range of adhesion proteins which may be essential for their uptake by endothelial cells (Fig. 3D). These adhesion proteins include PECAM1, MCAM, as well as integrins α5 and β1, which were approximately 3 to 9 times more abundant in LSS-lEVs. Interestingly, we found that endolysosomal and mitochondrial proteins were also more enriched in LSS-lEVS, which may be a sign of increased secretion of digestion products. Thus, quantitative proteomic analysis demonstrated that shear stress drastically alters the protein content of endothelial EVs.

### LSS-lEVs are enriched in adhesion proteins

We then sought to confirm the relative abundance of candidates identified by our proteomic analysis. We evaluated the levels of the different adhesion proteins in both lEVs and their parental cells by Western blot. We found that shear stress did not affect the cellular levels of most of the candidates, except for the long isoform of MCAM, which was more abundant in HUVECs exposed to LSS conditions (Fig. 4). However, PECAM-1, MCAM (long isoform), as well as integrins α5, α6 and β1 were found to be enriched in LSS-lEVs compared to HSS-lEVs, thereby confirming the results obtained by quantitative proteomics.

**Figure 4:**
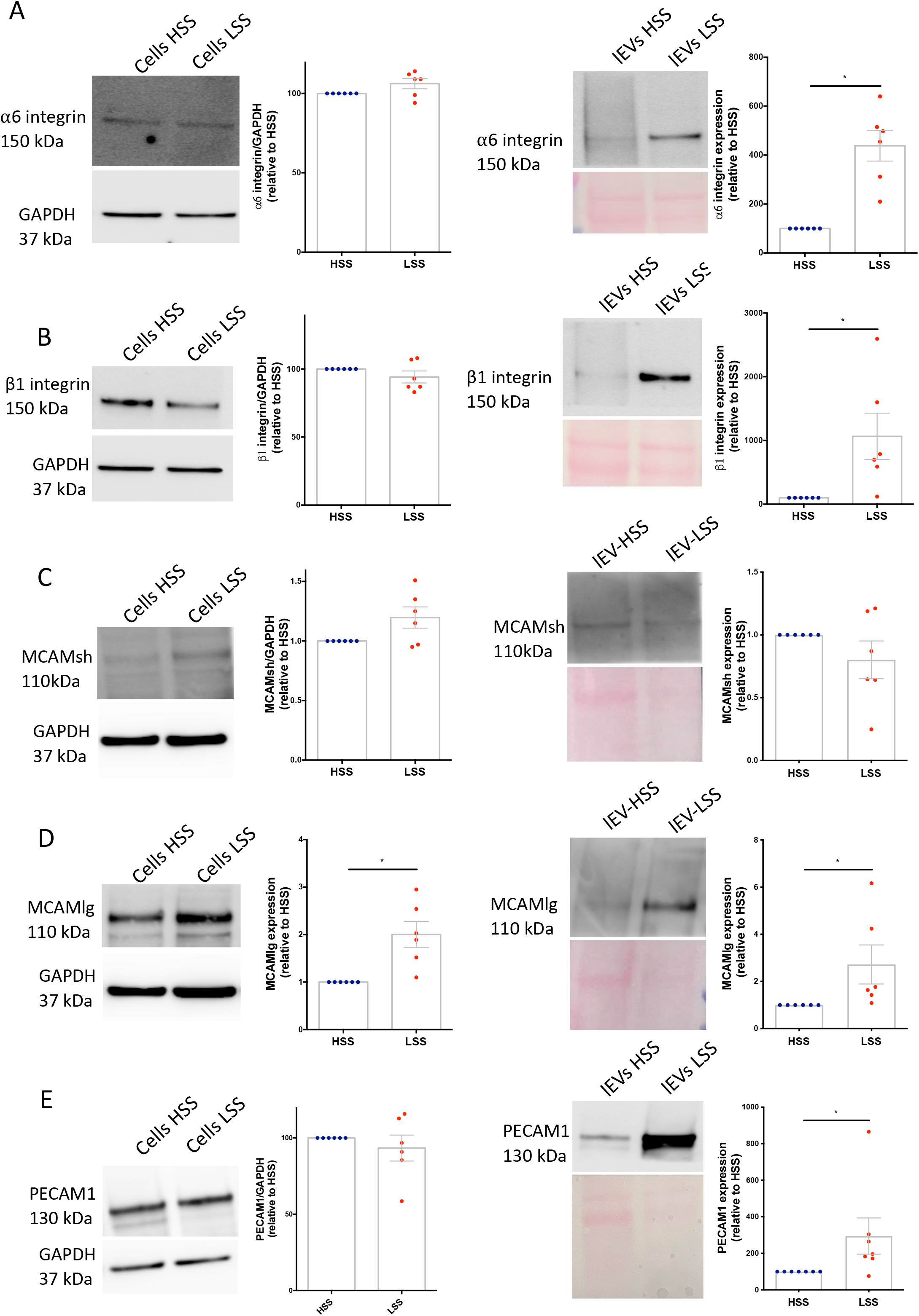
Adhesion proteins are more abundant in LSS-lEVs. Western blot analysis of α5 integrin **(A)**, β1 integrin **(B)**, MCAM short isoform **(C)**, MCAM long isoform **(D)**, and PECAM1 **(E)** levels in HUVEC cellular (left panels) and lEV (right panels) protein lysates. Data represent means ± SEM of 6 independent experiments, *P<0.05, Wilcoxon.

We next assessed the presence of the different adhesion proteins at the surface of lEVs by sensitive flow cytometry. lEVs produced by HUVECs exposed to HSS or LSS conditions were first stained with Membright so as to discern them from background noise. We then stained EV surfaces with antibodies directed against PECAM-1, MCAM, and integrins α5, α6 and β1. All five of these candidates were significantly enriched at the surface of LSS-lEVs when compared to HSS-lEVs (Fig. 5). Triton detergent exposure decreased over 90% of the positive labeling (Supp. Fig. 2).

**Figure 5:**
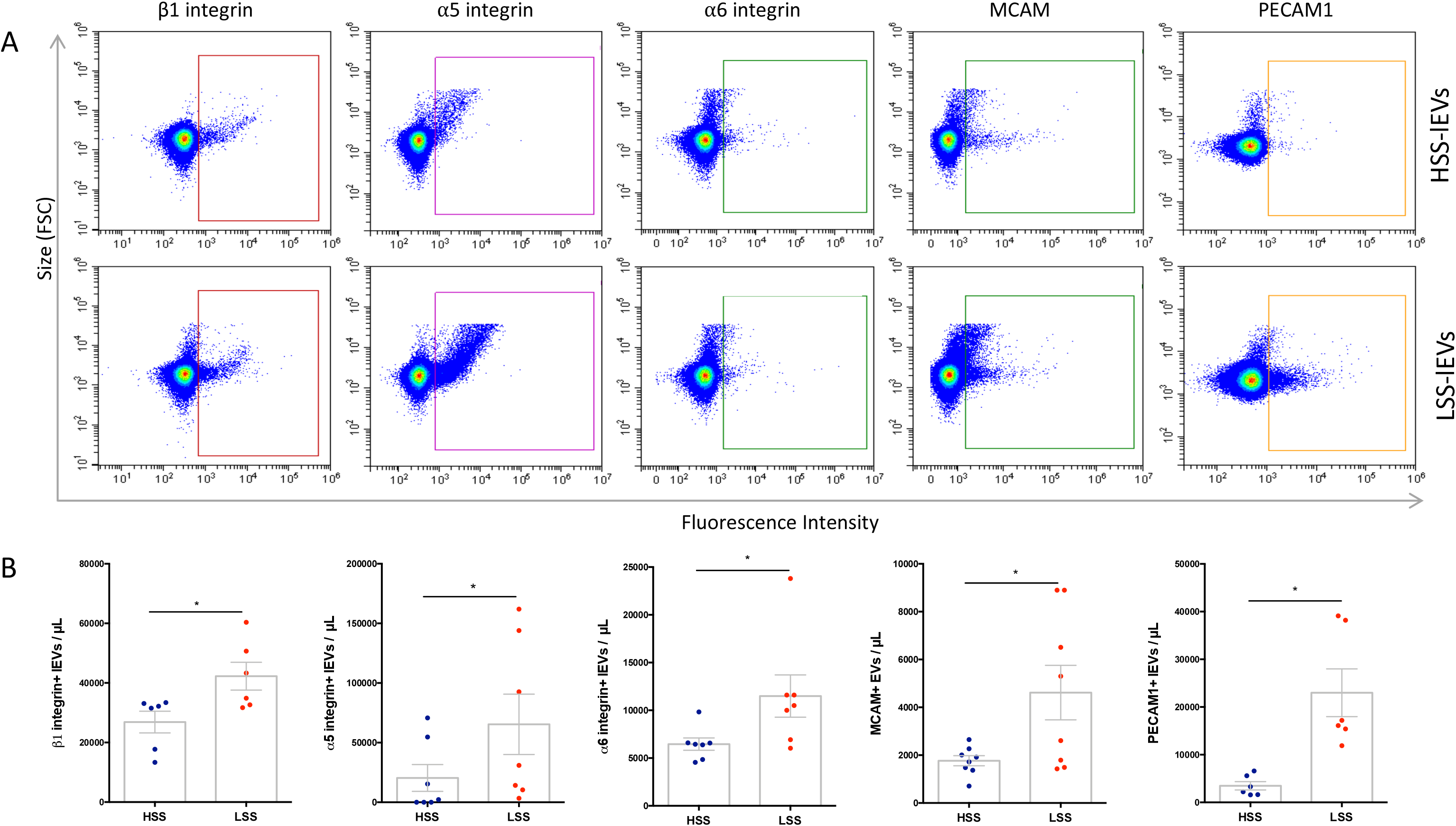
Adhesion protein are enriched at the surface of LSS-lEVs. **(A)** Representative flow cytometry dot plots of HUVEC HSS-lEVs (top panels) or LSS-lEVs (bottom panels) labeled with antibodies against β1 integrin, α5 integrin, α6 integrin, MCAM or PECAM1. **(B)** Data represent means ± SEM of 6-8 independent experiments. *P<0.05, Wilcoxon.

### LSS-lEV uptake by HUVECs is in part mediated by PECAM-1 and MCAM

Having identified the several adhesion proteins enriched at the surface of LSS-lEVs, we wanted to verify is they were functional and involved in the uptake process by endothelial cells. We assessed the potential role of PECAM-1, MCAM and integrins α5, α6 and β1 using neutralizing antibodies. Pre-incubating lEVs with anti-MCAM and anti-PECAM-1 antibodies resulted in a reduction of their uptake in a dose dependent manner (Supp. Fig. 3A). At a concentration of 10 μg/mL, anti-MCAM and anti-PEACM-1 neutralizing antibodies blocked uptake by approximately 40% and 30% respectively, while blocking the different integrins had no impact on lEV uptake by endothelial cells (Fig. 6A). Surprisingly, there was no additive effect of neutralizing both PECAM1 and MCAM (Supp. Fig. 3B). We also assessed the potential role of these adhesion proteins in the uptake of endothelial EVs by monocytic cells (THP-1). Here, we found that blocking PECAM-1 and integrins α5, α6 or β1 significantly reduced uptake, while blocking MCAM had no effect (Fig. 6B). These data suggest that the surface expression of PECAM-1 and MCAM is partially responsible for the uptake of LSS-lEVs by HUVECs, while the presence of integrins may be useful for incorporation by other cell types such as monocytes.

**Figure 6:**
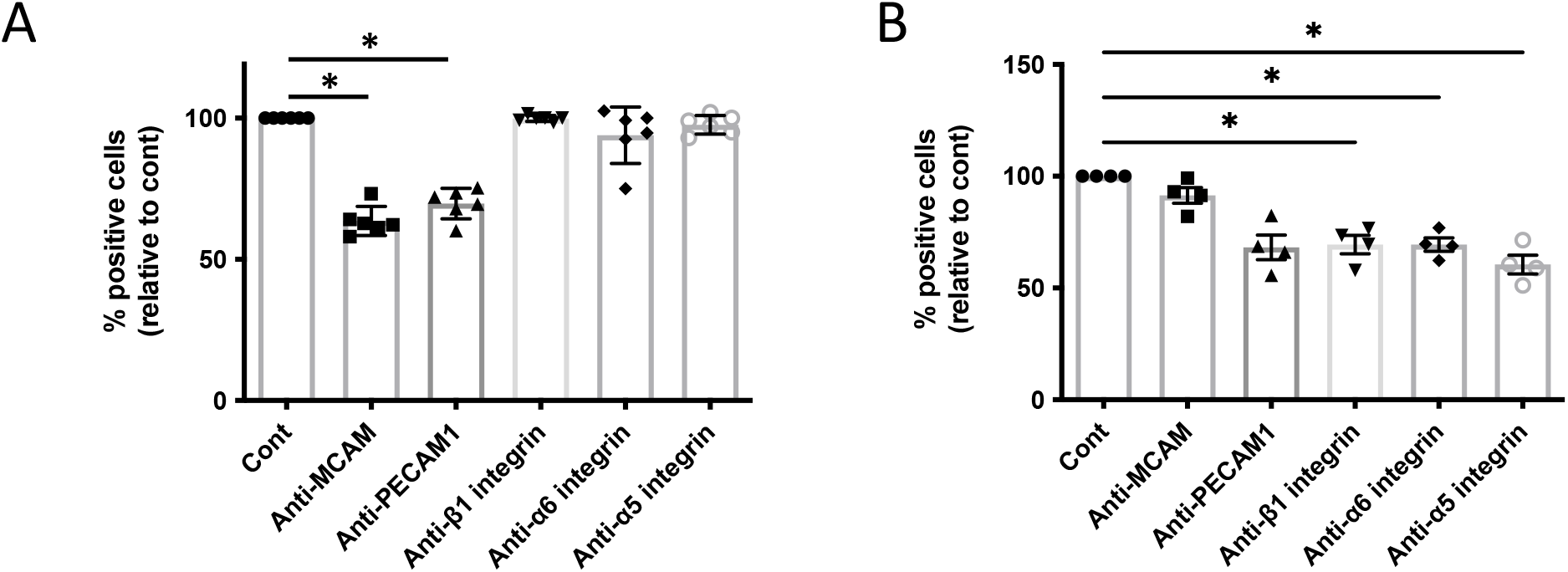
MCAM and PECAM1 are involved in endothelia EV uptake by HUVECs. Fluorescently-labeled HUVEC-derived LSS-lEVs were pre-incubated with neutralizing antibodies targeting MCAM, PECAM1, β1 integrin, α5 integrin or α6 integrin. lEVs were then incubated for 90 min with HUVECs exposed to LSS conditions for 24 h **(A)** or THP1 cells **(B)**. % of cells positive for EV signal, relative to control lEVs (Cont) was analyzed by flow cytometry. Data represent means ± SEM of 4-6 independent experiments. *P<0.05, Friedman test.

### Exposing endothelial cells to LSS increases the release of mitochondrial and endolysosomal material in lEVs

As our proteomic study had revealed an enrichment of mitochondrial and endolysosomal proteins in LSS-lEVs, we sought to validate this by Western blot and sensitive flow cytometry. While there was no difference the cellular levels the mitochondrial protein Tomm20 or the endolysosomal protein LAMP1 (Fig. 7A and 7C), we found higher levels of both markers in lEVs produced in LSS conditions (Fig. 7B and 7D). Similarly, LSS-lEV showed a higher staining for Mitotracker and Lysotracker (Fig. 7E and 7F). These data suggest that endothelial cells exposed to LSS redirect mitochondrial and endolysosomal material towards a secretory pathway.

**Figure 7:**
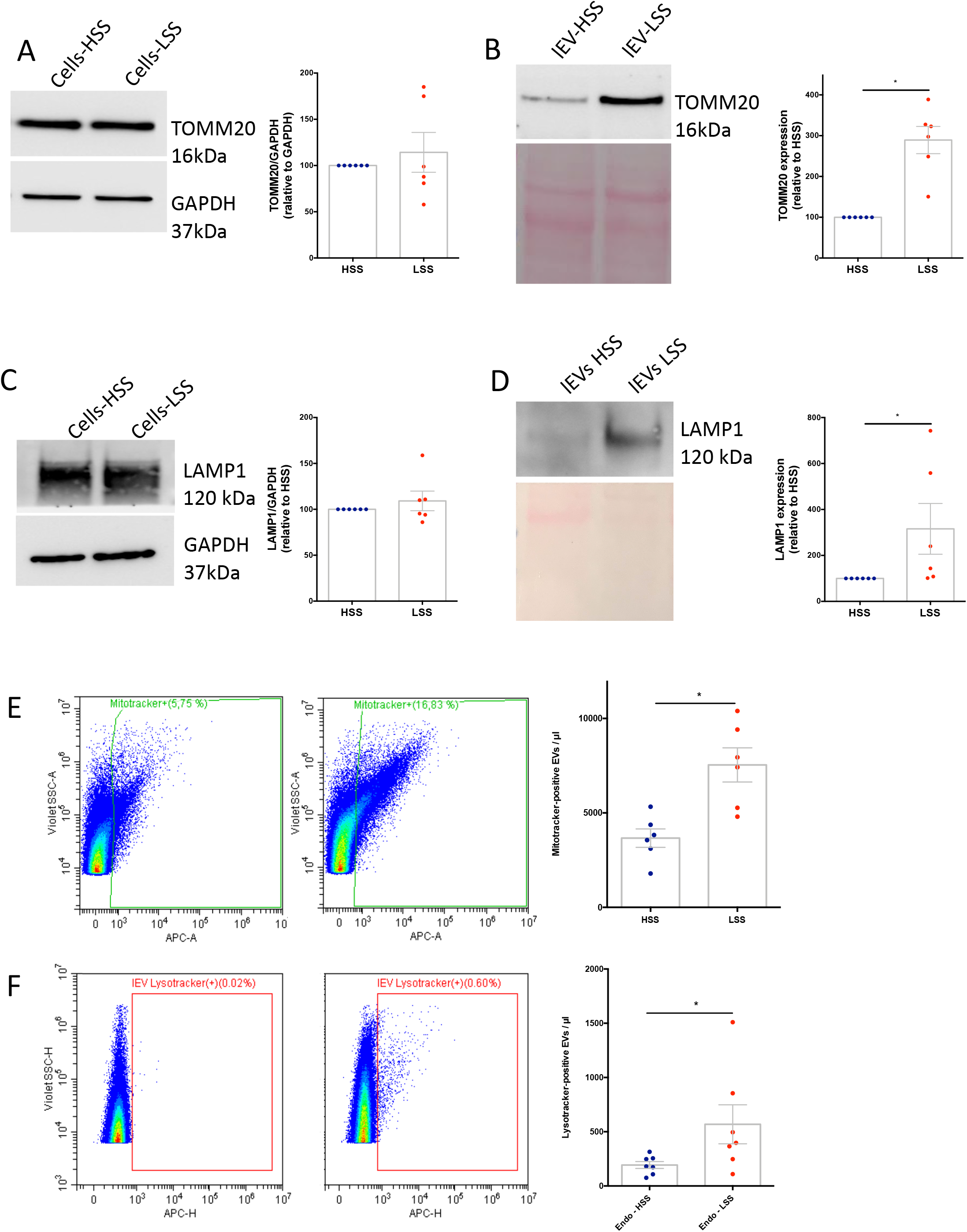
LSS-lEVs are enriched in mitochondrial and endolysosomal material. Western blot analysis of TOMM20 **(A and B)** and LAMP1 **(C and D)** levels in HUVEC cellular (left panels) and lEV (right panels) protein lysates. HSS-lEV and LSS-lEV were stained with Mitotracker **(E)** or Lysotracker **(F)** and analyzed by flow cytometry. Data represent means ± SEM of 6-7 independent experiments, *P<0.05, Wilcoxon.

### Uptake of LSS-lEVs elevates ROS levels in endothelial cells

We next wondered if the EV content could be transferred to recipient cells, and how that might affect their phenotype. Prestaining EVs with the Mitotracker dye allowed us to follow the transfer of mitochondrial material from EVs to cells. Cells exposed to LSS-lEVs presented higher Mitotracker signal than those exposed to HSS-lEVs, indicating higher incorporation of mitochondrial material (Fig. 8A). Similarly, confocal imaging revealed higher staining in cells exposed to LSS lEVs (Fig 8B).

**Figure 8:**
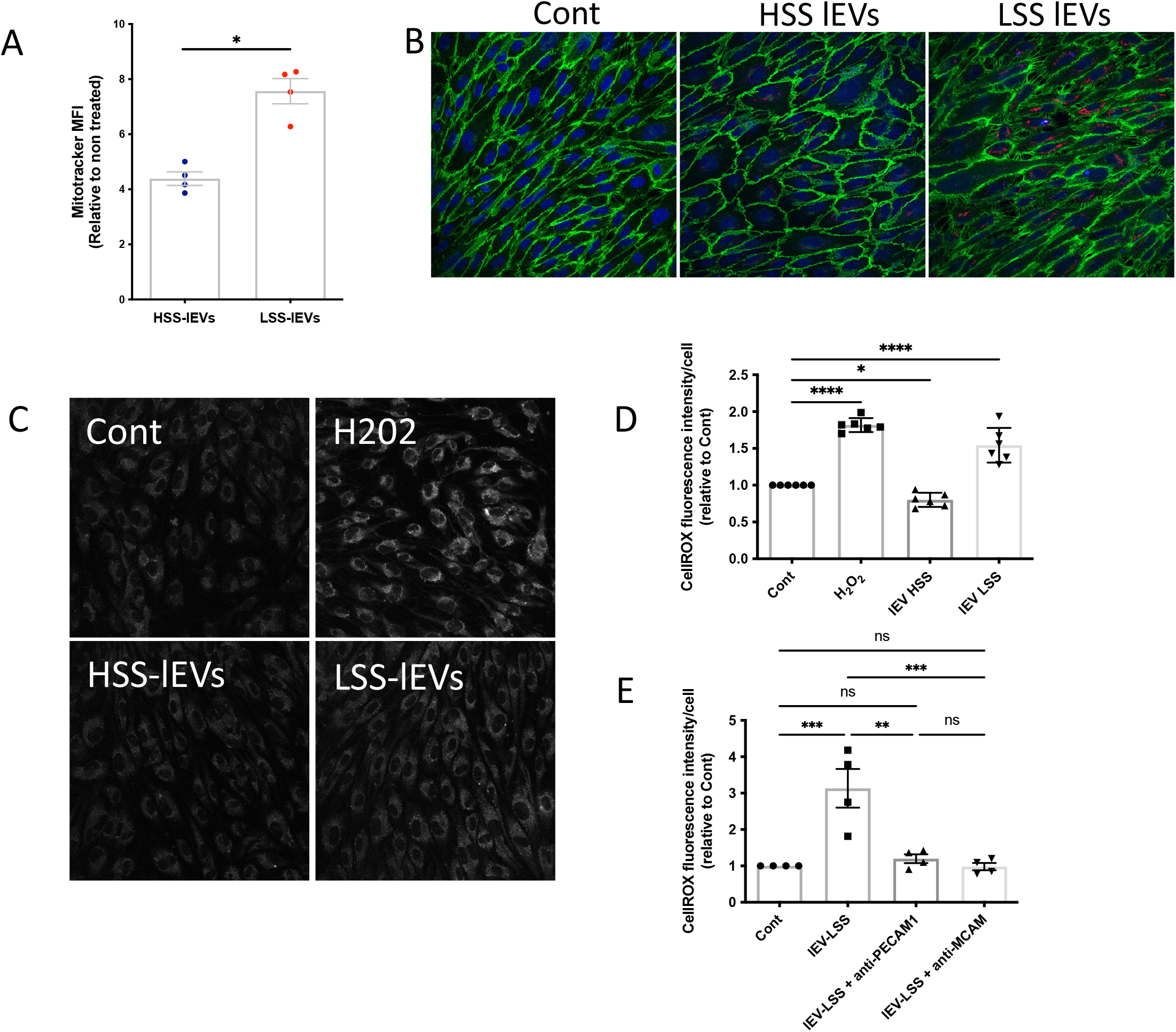
Effect of LSS-lEVs on endothelial ROS levels. **(A)** LSS-lEVs were pre-stained with Mitotracker and deposited on HUVECs for 4 h. Mitotracker median fluorescent intensity per cell, relative to cells treated without EVs, was analyzed by flow cytometry. Data represent means ± SEM of 4 independent experiments, *P<0.05, Paired t test. **(B)** LSS-lEVs were pre-stained with Mitotracker and deposited on HUVECs for 4 h. Cells were then stained for Cadherin-5 (green) and imaged by confocal microscopy. **(C and D)** HUVECs were incubated overnight with or without HSS-lEVs or LSS-lEVs. H_2_O_2_ was added in corresponding condition for the last hour of the experiment. Cells were stained with CellROX and immediately imaged. Fluorescent signal intensity was quantified from confocal images. Data represent means of fluorescence intensity per cell, relative to non-treated cells (Cont), ± SEM of 6 independent experiments. **(E)** lEVs were pre-incubated with or without neutralizing antibodies against PECAM1 or MCAM. lEVs were then incubated overnight with HUVECs. Cells were stained with CellROX and immediately imaged. Fluorescent signal intensity was quantified from confocal images. Data represent means of fluorescence intensity per cell, relative to non-treated cells (Cont), ± SEM of 4 independent experiments. *P<0.05, **P<0.01, ***P<0.001, ****P<0.0001, Friedman test.

As damaged mitochondria are a known source of reactive oxygen species (ROS; Zorov et al., 2014), we investigated ROS levels in HUVECs exposed to lEVs produced under HSS or LSS conditions. We used a 30 min hydrogen peroxide treatment as a positive control and found that it almost doubled ROS levels in HUVECs. Overnight exposure of endothelial cells to HSS-lEVs resulted in a 20% decrease in ROS levels. Conversely, exposing cells to LSS-lEVs led to 50% increase in ROS levels (Fig. 8C and 8D). Interestingly, the pro-oxidative effects of LSS-lEVs were completely blocked when EVs were pre-incubated with PECAM1- or MCAM-neutralizing antibodies (Fig. 8E). This indicates that the pro-oxidative effects are caused by adhesion protein-mediated attachment and/or endocytosis of EVs.

## Discussion

Over the past decade, EVs have emerged as crucial intercellular communication vehicles. By lining the vessel wall, endothelial cells are ideally positioned to be a major contributor to circulating EVs. Endothelial-EV composition and uptake is influenced by multiple factors including inflammation and thrombosis (Mathiesen et al., 2021). In this study, we demonstrate that shear stress is a key determinant in the way endothelial cell both produce and uptake EVs.

Our first major finding was that there was a higher concentration of EVs in the conditioned media of HUVECs cultured under HSS conditions compared to LSS conditions. As the number of EVs in the extracellular space is a balance between their rate of release and uptake, it is difficult at this point to conclude on the relative impact of each factor. In this study, however we were able to show, *in vitro* and *in vivo*, that endothelial cells exposed to LSS conditions exhibited a higher rate of EV uptake than EVs exposed to HSS, mainly through a clathrin-dependent pathway. This was observed for EVs of different cellular origin, including platelets, red blood cells, or peripheral blood mononuclear cells, but the most robust uptake was observed for endothelial EVs released under low shear stress (LSS-lEVs). Altogether, the present data support the hypothesis that endothelial EVs present in the blood can be re-incorporated by endothelial cells of the arteries. Using imaging flow cytometry, Banizs et al. had also observed that EV uptake by mouse aortic endothelial cells was essentially a receptor-mediated clathrin-dependent process. Higher endocytic activity under low or oscillatory shear stress does not seem to be restricted to endothelial EVs however, as it has also been described for erythrocyte derived EVs, as well as polystyrene or gold nanoparticles (Charwat et al., 2018; Qin et al., 2021). This suggests that the uptake of various types of particles is enhanced by low magnitude shear stress, and implies that atheroprone regions of the arterial tree may be targetable by therapeutic nanovesicles. While the underlying mechanisms remain to be elucidated, Qin et al. suggest that LSS-induced oxidative stress may be implicated, as oxidative stress genes are expressed earlier that pro-inflammatory genes in response to LSS, and antioxidant treatment reduced EV uptake (Ajami et al., 2017; de Keulenaer et al., 1998; Hsiai et al., 2007; Hsieh et al., 2014).

The uptake mechanism involves protein interactions which facilitate EV retention and subsequent endocytosis. Our data showed that LSS-lEVs were more adherent than HSS-lEVs, we therefore sought to identify adhesion proteins that could mediate these effects. Using proteomic analysis, combined with Western blot and sensitive flow cytometry, we were able to uncover several adhesion proteins that are enriched at the surface of LSS-lEVs. Since most of these proteins showed no significant differences in cellular proteins levels, we hypothesize that they were selectively directed towards a secretory pathway in LSS conditions. As key players in cellular adhesion, several integrins have been shown to be crucial in cell-EV interactions (Deregibus et al., 2007; Esmaeili et al., 2022; Mulcahy et al., 2014; Rana et al., 2012). Here we found that integrins α5, α6 and β1 were enriched at the surface of LSS-lEVs. Though they were involved in endothelial EV uptake by monocytes, we found that they played little role in uptake by endothelial cells. PECAM1 and MCAM, on the other hand, were required for EV uptake by endothelial cells. While MCAM had previously been implicated in the interaction of melanoma cell-derived EVs with endothelial cells (Ghoroghi et al., 2021), this represents, to our knowledge, the first description of an involvement of PECAM1 in EV uptake by endothelial cells. Interestingly, there was no additive effect of neutralizing both MCAM and PECAM1, suggesting that they may be located on the same subset of EVs, and may work in tandem to allow EV endocytosis. Together, these data reinforce the idea that EV internalization is a selective process. Surface proteins seem to serve as molecular barcodes, targeting EVs towards specific recipient cells, and therefore playing a role in the biodistribution of circulating EVs.

Several studies have shown evidence for the presence of extracellular mitochondria, either freely (al Amir Dache et al., 2020; Joshi et al., 2019; Puhm et al., 2019) or encapsulated within EVs (D’Acunzo et al., 2021; Todkar et al., 2021; Torralba et al., 2016). Although the intracellular mechanisms regulating this process are still emerging, evidence points towards the involvement of mitochondria-derived vesicles. These are small vesicles which deliver damaged mitochondrial content to late endosomes/lysosomes in a Pink1/Parkin-dependent manner (Sugiura et al., 2014). Remarkably, autophagy-deficient cells displayed higher extracellular mitochondria release, implying that perturbation of the mitophagy pathway prompts mitochondria expulsion (Choong et al., 2021). Thus, this process appears to be a comparable yet distinct quality control pathway from conventional mitophagy. It might be involved in ensuring mitochondrial homeostasis in the event of mild mitochondrial damage, such as oxidative stress, while mitophagy is triggered by acute mitochondrial depolarization (Amari & Germain, 2021). Our proteomic analysis of endothelial EVs revealed an enrichment of endolysosomal and mitochondrial proteins in LSS-lEVs. As we had previously demonstrated that the autophagic flux is hindered in LSS conditions (Vion et al., 2017), we hypothesize that endothelial cells respond by packaging damaged mitochondria for secretion in EVs.

Our data indicate that this mitochondrial material can be transferred to new endothelial cells via EVs, and may be involved in elevating ROS levels in recipient cells. This pro-oxidative effect was blocked by neutralizing antibodies targeting PECAM1 or MCAM, which suggests that it is mediated by EV uptake. As mentioned above, endothelial cells exposed to LSS seem to have increased endocytic activity but reduced degradative capacity. Together, these processes may lead to accumulation of noxious pro-oxidative material which contributes to sensitizing cells to inflammatory stimuli in atheroprone regions (Puhm et al., 2019; Zorov et al., 2014). Interestingly, exposure to HSS-lEVs had a small antioxidant effect, even though these EVs were barely taken up. Notably, we were able to detect several components of the anti-oxidant machinery in HSS-lEVs, including Glutathione S transferase, Peroxiredoxin-1/2/6 and Glutamate dehydrogenase 1. While their eventual implication in these effects remains to be determined, previous work has shown a transfer of anti-oxidant machinery via EVs (Bodega et al., 2017, 2018). It is therefore tempting to imagine leveraging the knowledge obtained in this study to target therapeutic material to endothelial cells in areas of LSS. In that sense. Engineering HSS-EVs to overexpress MCAM and/or PECAM1 could bolster their uptake and anti-oxidant effects on recipient cells.

In conclusion, endothelial cells exposed to atheroprone LSS are characterized by higher incorporation of circulating endothelial EVs than those exposed to HSS. LSS endothelial cells also release EVs enriched in adhesion proteins and mitochondrial material, which may transfer noxious pro-oxidative cargos to recipient cells.

## Materials and Methods

### Cell culture

Confluent Human Umbilical Vein Endothelial Cells (HUVEC; passage 2-4; 20 different primary cultures; PromoCell) were cultured on 0,2 % gelatin-coated slides, in Endothelial Cell Basal Medium (ECBM, PromoCell), supplemented with 2 % Fetal Calf Serum (PromoCell), growth factors (0.4% ECGS, 0.1 ng/mL EGF, 1 ng/mL ß-FGF), heparin (90 μg/mL), hydrocortisone (1 μg/mL), Amphotericin B (10 μg/L, Gibco), Streptomycin (100 IU/mL, Gibco) and Penicillin (100 IU/mL, Gibco). Murine endothelial cells, SVEC4-10 (ATCC), were cultured in DMEM supplemented with Fetal Calf Serum (10 %), Amphotericin B (10 μg/L), Streptomycin (100 IU/mL) and Penicillin (100 IU/mL, Gibco). The human monocytic cell line, THP-1 (ATCC), was maintained in RPMI 1640 medium supplemented with Fetal Calf Serum (10 %), Amphotericin B (10 μg/L), Streptomycin (100 IU/mL) and Penicillin (100 IU/mL, Gibco).

### *In Vitro* shear stress system

A controlled level of laminar shear stress was applied to confluent endothelial cells using a parallel plate chamber connected to a perfusion circuit driven by a peristaltic pump (Gilson). Before use, the fetal calf serum was ultracentrifuged (100,000 x *g*, 90 min) and the complete medium was filtered on a 0.1 μm membrane to remove serum EVs. Cells were placed in the perfusion system for 24 hours, in a sterile 5 % CO2 incubator at 37°C. LSS (2 dyn/cm^2^) or HSS (20 dyn/cm^2^ for HUVECs and 40 dyn/cm^2^ for SVEC4-10) was calculated using Poiseuille’s law.

### Endothelial EV isolation

Following exposure to shear stress, the conditioned media were collected and first centrifuged at 600 x *g* for 15 min at 4°C to remove cell debris, then at 4,500 x *g* for 20 min at 4°C to remove apoptotic bodies. Resulting supernatants was subjected to differential centrifugations at 20,500 x *g* to pellet lEVs and 100,000 x *g* to pellet sEVs. EVs were resuspended in filtered PBS, purified by size exclusion chromatography (Izon, New Zealand), and stored at -80°C until use. Blood was collected in citrated tubes. Tubes were centrifuged at 2500 x *g* for 15 min. The platelet-free plasma supernatant (PFP) was collected and subjected to size exclusion chromatography to isolate EVs. Blood cells (platelets, neutrophils, peripheral blood mononuclear cells, and red blood cells) were separated on a density gradient (Granulosep, Eurobio) *via* a 30 min centrifugation at 700 x *g*. EVs were obtained by incubating red blood cell with CaCl (5 mM) and other cell types with the calcium ionophore A23187 (1 mM, C9275, Sigma) for 30 min at 37°C. Samples were then centrifuged at 600 x *g* for 10 min and 4,500 x *g* for 20 min to eliminate cells and debris. lEVs were obtained as described above.

### Transmission electron microscopy

EVs were directly deposited on formvar/carbon coated copper/palladium grids for 20 min at room temperature and fixed with PFA 2 % / 0.1 M phosphate buffer. Negative staining was performed using uranyl acetate 0.4 % in methylcellulose. The samples were analyzed with a 80 kV transmission electron microscope (Tecnai Spirit G2; ThermoFischer Scientific) equipped with a 4k CCD camera (On view 4k x 4k Gatan).

### Tunable Resistive Pulse Sensing Measurements

EVs were quantified using NP400 (lEVs) and NP150 (sEVs) nanopores (Izon, New Zealand). The measurements were calibrated with polystyrene beads CPC 400 (350 nm mode diameter) and CPC 200 (210 nm mode diameter) respectively. Concentrations were expressed as EVs normalized to the volume of conditioned medium collected after exposure to shear stress.

### EV Flow Cytometry

lEVs were incubated with the membrane probe MemBright (1 μM, Idylle Labs) for 30 minutes at room temperature. They were then rinsed in 1 mL of PBS and pelleted by centrifugation. EVs were resuspended in PBS and used immediately for flow cytometry experiments. lEVs were analysed on a Cytoflex flow cytometer (Beckman Coulter, USA) using Megamix-Plus SSC beads and Megamix-Plus SSC beads (Biocytex, France) to define events of 0.1 to 0.9 μm diameter size. For experiments using fluorescence-conjugated antibodies to stain lEV surface markers, antibodies were first centrifuged for 5 min at 13,000 × g at 4°C before being added to EV samples. lEVs were defined as events stained by MemBright. Antibodies were as follows: PECAM1 (IM2409, Beckman Coulter), MCAM (A07483, Beckman Coulter), α5 integrin (563578, BD Biosciences), α6 integrin (561894, BD Biosciences) and β1 integrin (130-101-271, Miltenyi Biotec). Corresponding isotype controls were as follows: IgG1 Mouse-PE (A0779, Beckman Coulter), IgG2a Mouse-PE (A09142, Beckman Coulter), IgG1κ Mouse-AF647 (565571 BD Biosciences), IgG2a κ Rat-PE (551799, BD Biosciences) and IgG1κ Mouse-APC (130-113-196, Miltenyi Biotec) respectively. To assess levels of mitochondrial and lysosomal material, EVs were stained with Mitotracker Deep Red (400 nM, ThermoFisher Scientific) and LysoTracker Deep Red (50 nM, ThermoFisher Scientific) respectively. Controls include detergent lysis, buffer-only, buffer with reagent (without EVs) and unstained samples.

### Immunoblotting

HUVECs were washed with cold PBS and lysed in Radioimmunoprecipitation assay (RIPA) buffer. For EV samples, RIPA (2X) was added to equal volume of EVs suspended in PBS. Protein content was quantified using the Micro BCA Protein Assay kit (ThermoFisher Scientific). Lysates were mixed with the reducing sample buffer for electrophoresis and subsequently transferred onto nitrocellulose membranes (Bio-Rad). Equal loading was checked using Ponceau Red solution. Membranes were incubated with primary antibodies overnight at 4°C, with constant agitation. After secondary antibody incubation (1 h, room temperature), immunodetection was performed using Clarity™ Western ECL Substrate, and the chemiluminescent signal was revealed using the LAS-4000 imaging system and quantified with MultiGauge software (Fujifilm, Japan). Antibodies were as follows: CD9 (Merck Millipore, CBL162), CD63 (MBL Life Science, D263-3), HSC70 (Enzo Life Sciences, SPA815), TOMM20 (Abcam, ab46798), LAMP1 (BD Biosciences 611043), GAPDH (Merk Millipore, mab-374), α6 integrin (Cell Signaling Technology, 3750), β1 integrin (Abcam, ab52971), MCAM long and short (generously provided by Dr Aurélie Leroyer, Aix-Marseille University), HRP-linked Anti-rabbit (GE Healthcare, NA934), HRP-linked Anti-mouse (GE Healthcare, NXA931), HRP-linked Anti-rat (Jackson Immunity, 112-035-150).

### Uptake assay by Flow cytometry

Following shear stress experiments, cells were washed with warm media and immediately incubated with Membright-labelled EVs (7.5 million/mL) for indicated periods. Cells were then washed with PBS, detached with Trypsin and the fluorescent signal intensity per cell was measured by flow cytometry. When indicated, cells were treated with amiloride (1 mM, Sigma, A7410), nystatin (50 μM, Sigma N1638) or chlorpromazine (20 μM, Sigma, C8138) 1 h before incubation with EVs. For surface protein neutralization, EVs were pre-incubated with blocking antibodies for 30 min at room temperature, then centrifuged to eliminate unbound antibodies. EVs were then resuspended in warm media and deposited on cells. Blocking antibodies were as follows: PECAM1 (MA3100, ThermoFisher Scientific), MCAM (clone S-Endo 1, BioCytex), β1 integrin (Biolegend, 921303), α5 integrin (Biolegend, 921704), α6 integrin (Biolegend, 313602).

### Immunofluorescent staining and microscopy *in vitro*

After shear stress, HUVECs were fixed with 4 % PFA and blocked with 5% BSA in PBS. Cells were incubated with primary antibody overnight at 4°C (Cadherin-5, Santa Cruz Biotechnology, sc-6458). And then with AlexaFluor-conjugated secondary antibody (ThermoFisher Scientific). Nuclei were stained with DAPI (0.1 μg/mL).

### ROS measurement

To assess ROS levels following exposure to EVs, we used the CellROX Deep Red Reagent (Thermo Fisher Scientific, C10422), a fluorogenic probe designed to reliably measure ROS inside living cells. This cell-permeable dye is in a non-fluorescent reduced state while outside the cell and exhibits excitation/emission maxima at 640/665 nm upon oxidation. HUVECs were incubated with 5 μM CellROX for 30 minutes at 37°C. Cells were the then fixed and imaged by confocal microscopy.

### *In vivo* biodistribution studies

Mice used in the study were of C57BL/6 genetic background. Membright-labeled SVEC4-10-derived EVs (500 x 10^6^ in 150 μl PBS) were injected in the retro-orbital sinus. PBS vehicle and EV-labeling solution supernatant was used as controls. Mice were sacrificed 30 min post-injection and their aortas were injected in situ with PBS supplemented with 4 % PFA. After dissection, the aortas were blocked with 5 % BSA for 20 min and stained with anti-Cadherin-5 antibody (Santa Cruz biotechnology, sc-6458) and DAPI. For all mice, 10 confocal images were obtained from regions in the aortic arch (LSS) and the thoracic aorta (HSS). All experiments were performed in accordance with the European Community guidelines for the care and use of laboratory animals (no. 07430) and were approved by the institutional ethical committee (no. 02526.02).

### Mass spectrometry analysis

Five independent sets of experiments were performed. Purified EVs were lysed in freshly prepared urea buffer (8M urea, 200 mM ammonium bicarbonate). For mass spectrometry analysis, proteins were precipitated overnight at –20°C with 0.1 mol/L ammonium acetate glacial in 80% methanol (buffer 1). After centrifugation at 14,000 g and 4°C for 15 minutes, the resulting pellets were washed twice with 100 μL buffer 1 and further dried under vacuum (Savant Centrifuge SpeedVac concentrator, Thermo Fisher Scientific). Proteins were then reduced by incubation with 10 μL of 5 mmol/L DTT at 57°C for 1 hour and alkylated with 2 μL of 55 mmol/L iodoacetamide for 30 minutes at room temperature in the dark. Trypsin/LysC (Promega) was added twice at 1:100 (wt/wt) enzyme/substrate, at 37°C for 2 hours first and then overnight. Samples were then loaded onto a homemade C18 StageTips for desalting. Peptides were eluted using 40:60 MeCN/H2O plus 0.1% formic acid and vacuum concentrated to dryness. Online chromatography was performed with an RSLCnano system (Ultimate 3000, Thermo Fisher Scientific) coupled online to a Q Exactive HF-X with a Nanospray Flex ion source (Thermo Fisher Scientific). Peptides were first trapped on a C18 column (75-μm inner diameter × 2 cm; nanoViper Acclaim PepMap 100, Thermo Fisher Scientific) with buffer A (2:98 MeCN/H2O in 0.1% formic acid) at a flow rate of 2.5 μL/min over 4 minutes. Separation was then performed on a 50 cm × 75 μm C18 column (nanoViper Acclaim PepMap RSLC, 2 μm, 100 Å) regulated to a temperature of 50°C with a linear gradient of 2%–30% buffer B (100% MeCN in 0.1% formic acid) at a flow rate of 300 nL/min over 91 minutes. MS full scans were performed in the ultrahigh-field Orbitrap mass analyzer in the m/z range of 375–1500 with a resolution of 120,000 at m/z 200. The 20 most intense ions were subjected to Orbitrap for further fragmentation via high-energy collision dissociation (HCD) activation and a resolution of 15,000, with the intensity threshold kept at 1.3 × 105. We selected ions with charge state from 2+ to 6+ for screening. Normalized collision energy (NCE) was set at 27 and a dynamic exclusion of 40 seconds.

For identification, data were searched against the *Homo Sapiens* (UP000005640) UniProt database using Sequest HT through proteome discoverer (version 2.2). Enzyme specificity was set to trypsin, and a maximum of 2 missed cleavage sites were allowed. Oxidized methionine and N-terminal acetylation were set as variable modifications. Maximum allowed mass deviation was set to 10 ppm for monoisotopic precursor ions and 0.02 Da for MS/MS peaks. The resulting files were further processed using myProMS (Poullet et al., 2007) v3.6. FDR calculation used Percolator (Spivak et al., 2009) and was set to 1% at the peptide level for the whole study. The label-free quantification was performed by peptide extracted ion chromatograms (XICs) computed with MassChroQ version 2.2 (Valot et al., 2011). For protein quantification, XICs from proteotypic peptides shared by compared conditions (TopN matching) with 2 missed cleavages were used. Median and scale normalization was applied on the total signal to correct the XICs for each biological replicate. To estimate the significance of the change in protein abundance, a linear model (adjusted on peptides and biological replicates) was performed, and P values were adjusted with a Benjamini-Hochberg FDR procedure with a control threshold set to 0.05. Proteins with at least 3 distinct peptides in five replicates, a 2-fold enrichment and an adjusted p-value ≤ 0.05 were considered significantly enriched in sample comparisons. Proteins selected with these criteria were further subjected to Gene Ontology (GO) cellular compartment analysis.

### Statistical analysis

Data are expressed as mean ± SEM for all experiments. Statistical analyses were performed using the GraphPad Prism 9 statistical package. Normality of the data was confirmed using Shapiro-Wilk test and accordingly different statistical tests were used as described in legends. For data that follow gaussian distribution, t test or one-way Anova were used. For data that do not follow gaussian distribution, Wilcoxon or Friedman tests were used. p-Values smaller than 0.05 were considered as statistically significant. *, P<0.05, **, P<0.01, ***, P<0.001, ****, P<0.0001.

## Acknowledgement

The authors acknowledge the Flow Cytometry Facility manager, Dr Camille Knosp, the members of the INSERM U970 Histology, Microscopy and ERI facilities, as well as the ImagoSeine core facility of the Institute Jacques Monod, member of the France BioImaging infrastructure (ANR-10-INBS-04) and GIS-IBiSA.

## Author contribution

PMC and SC designed, performed, and analyzed most of the experiments with the help of KEJ, CD, and TN. FM performed and analyzed EV flow cytometry. MLC performed and analyzed electron microscopy. FD and DL performed and analyzed mass spectrometry. PMC, CMB and GvN wrote/reviewed the manuscript. CMB conceived and supervised the study.

## Funding

This work has been supported by INSERM and a grant from the French National Agency for Research ANR-16-CE14-0015-01 and from the Fondation pour la Recherche Médicale (FRM EQU202003010767). SC was supported by the USPC Inspire Program European Union’s Horizon 2020 research and innovation program under the Marie Skłodowska-Curie grant agreement No 665850. MLC was supported by the French National Agency for Research ANR-20-CE18-0026-01. PMC was partly supported by INCA-PLBIO19-059.

## Conflict of interest

All authors declare nothing to disclose

## Abbreviations

EVs: Extracellular Vesicles
HUVEC: Human Umbilical Vein Endothelial Cells
HSS: High Shear Stress
LSS: Low Shear Stress
MCAM: Melanoma Cell Adhesion Molecule
PECAM1: Platelet endothelial cell adhesion molecule 1

## Notes

### Competing Interest Statement

The authors have declared no competing interest.

