## Supplemental figures for "Atheroprone shear stress stimulates noxious endothelial extracellular vesicle uptake by MCAM and PECAM-1 cell adhesion molecules"

### Supplementary Figures

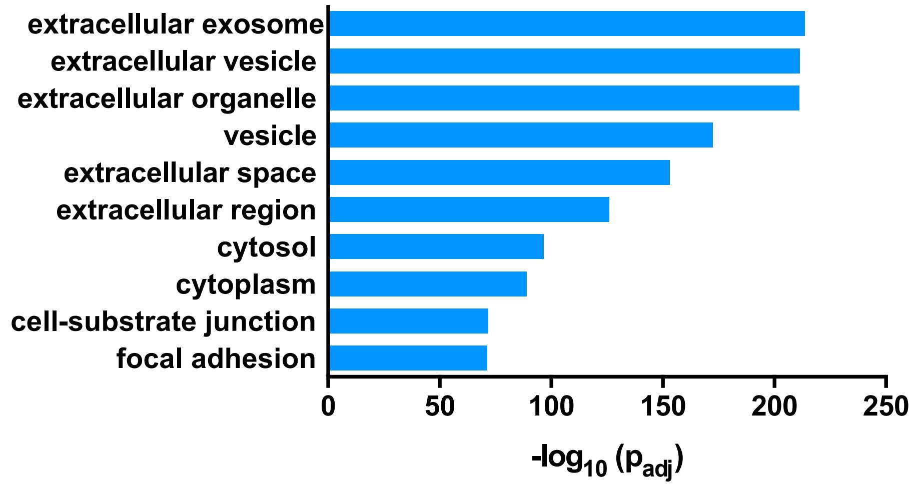


**Supplementary Figure 1:** Gene Ontology, cellular component terms associated with proteins identified in endothelial EVs.

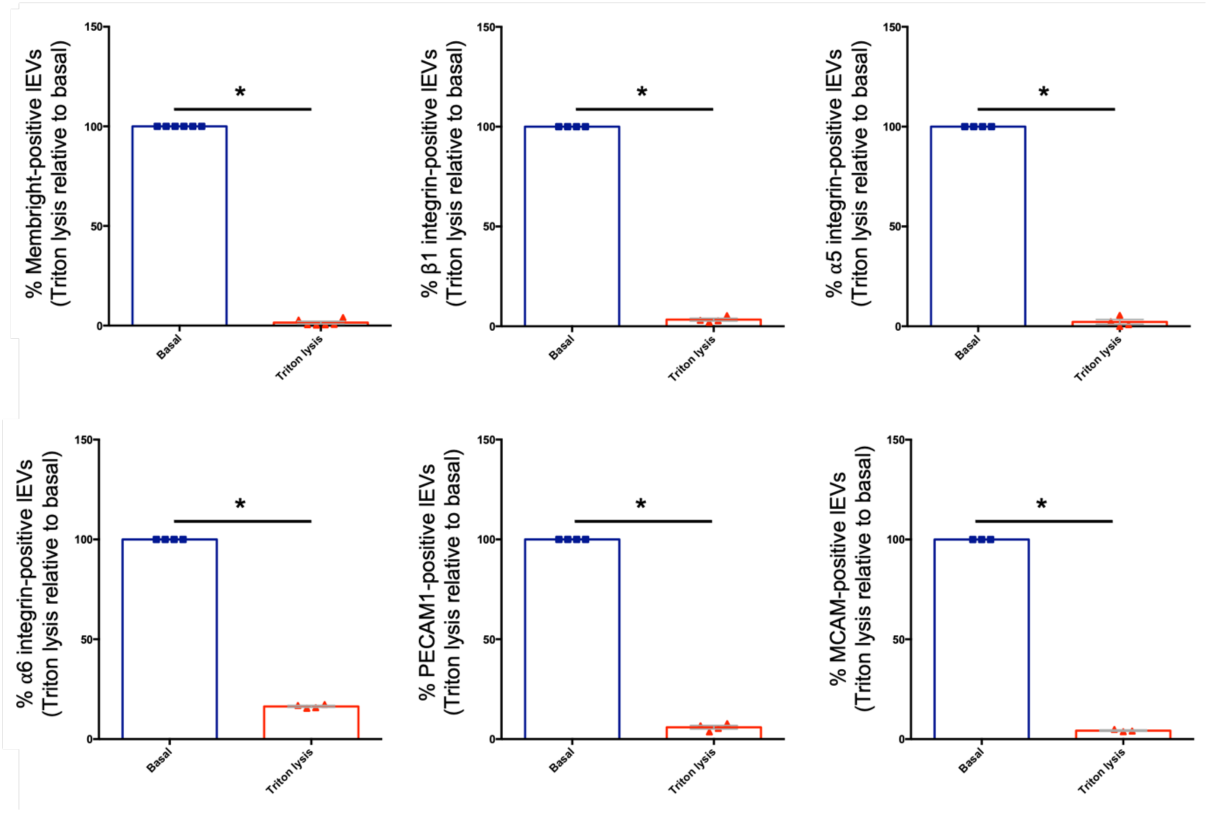


**Supplementary Figure 2**: Effect of Triton lysis on lEVs flow cytometry analysis. **(A)** Levels of lEVs positive for the different markers in absence or presence of Triton. Data are expressed as mean +/- SEM. *P < 0.05, paired t test.

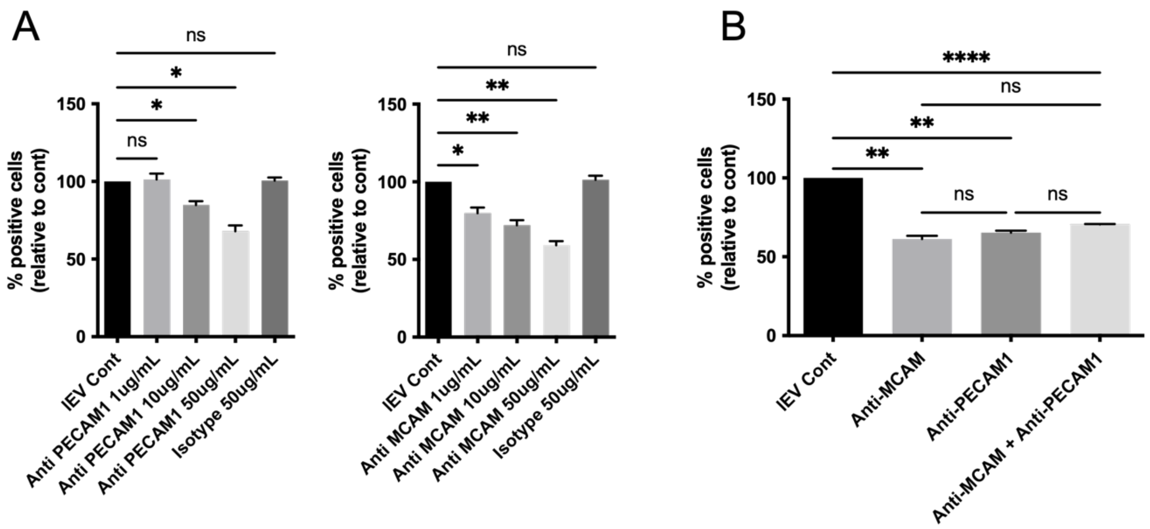


**Supplementary Figure 3:** Fluorescently-labeled HUVEC-derived LSS-lEVs were pre-incubated with different concentrations of either MCAM- or PECAM1-neutralizing antibodies **(A)** or with a combination of both **(B)**. lEVs were then incubated for 90 min with HUVECs. % of cells positive for EV signal, relative to control lEVs (Cont) was analyzed by flow cytometry. Data represent means ± SEM of 6-8 independent experiments. *P < 0.05, **P<0.01, ****P<0.0001; Friedman test.
